# Coatomer complex I is required for the transport of SARS-CoV-2 progeny virions from the endoplasmic reticulum-Golgi intermediate compartment

**DOI:** 10.1101/2024.04.10.588984

**Authors:** Ai Hirabayashi, Yukiko Muramoto, Toru Takenaga, Yugo Tsunoda, Mayumi Wakazaki, Mayuko Sato, Yoko Fujita-Fujiharu, Norimichi Nomura, Koji Yamauchi, Chiho Onishi, Masahiro Nakano, Kiminori Toyooka, Takeshi Noda

## Abstract

SARS-CoV-2 undergoes budding within the lumen of the endoplasmic reticulum-Golgi intermediate compartment (ERGIC) and delivers progeny virions to the cell surface by employing vesicular transport. However, the molecular mechanisms remain poorly understood. Using three-dimensional electron microscopic analysis, such as array tomography and electron tomography, we found that virion-transporting vesicles possessed a coated protein on their membrane and demonstrated that the coated protein was coatomer complex I (COPI). During the later stages of SARS-CoV-2 infection, we observed a notable alteration in the distribution of COPI and ERGIC throughout the cytoplasm. Depletion of COPB2, a key component of COPI, led to the confinement of SARS-CoV-2 structural proteins in the perinuclear region, where progeny virions were accumulated within the ERGIC. While the expression levels of viral proteins within cells were comparable, this depletion significantly reduced the efficiency of virion release, leading to the significant inhibition of viral replication. Hence, our findings suggest COPI as a critical player in facilitating the transport of SARS-CoV-2 progeny virions from the ERGIC. Thus, COPI could be a promising target for the development of antivirals against SARS-CoV-2.

## Introduction

Severe acute respiratory syndrome-coronavirus-2 (SARS-CoV-2), a member of the β-coronavirus family within *Coronaviridae*, is an enveloped RNA virus with a positive-stranded genome. It first emerged in late 2019 and has since spread rapidly worldwide, giving rise to various variants that adapt to human hosts and evade the immune system. Despite the development of effective vaccines that mitigate disease severity, coronavirus disease 2019 (COVID-19) continues to pose a significant threat to public health.

SARS-CoV-2 gains entry into host cells through the interaction between its spike protein (S) and the angiotensin-converting enzyme 2 (ACE2) receptor (1). Subsequent to the fusion of the viral envelope with either the plasma membrane or the endosomal membrane, following endocytosis (2), the positive-stranded RNA genome is released into the cytoplasm, initiating the synthesis of the RNA-dependent RNA complex (RdRp). RdRp transcribes negative-strand antigenomic RNAs, serving as templates for viral RNA genome replication and the production of sub-genomic mRNAs encoding viral proteins, such as S, membrane protein (M), envelope protein (E), and nucleocapsid protein (N), which collectively constitute the SARS-CoV-2 virus particles. Non-structural proteins 3, 4, and 6 (nsp3, nsp4, and nsp6, respectively) induce the formation of double-membrane vesicles (DMVs) (3–5), acting as sites for viral genome transcription and replication (6–8). Once all viral proteins are synthesized, viral assembly and the budding of progeny virions occur within the endoplasmic reticulum-Golgi intermediate compartment (ERGIC), and these progeny virions are transported to the cell surface via vesicular transport and the biosynthetic secretory pathways (6, 9, 10). Recently, Ghosh *et al.* reported that β-coronaviruses, including SARS-CoV-2, employ lysosomes for virion transport instead of the biosynthetic secretory pathway (11). Mendonca *et al.* demonstrated that virions exit through tunnels connecting unidentified large virion-rich vesicles to the plasma membrane using cryogenic focused ion beam-scanning electron microscopy (cryo FIB-SEM) (12). Furthermore, Eymieux *et al.* employed serial-ultrathin section transmission electron microscopy (TEM) to show that virions are primarily transported and released through small secretory vesicles containing a single virion rather than large virion-rich vesicles (13). While these findings are not mutually exclusive, the process of intracellular virion transport and the specific host factors involved in this process remain a subject of controversy and incomplete understanding.

To elucidate the intracellular transport mechanism of SARS-CoV-2 virions for their release, we initiated a comprehensive investigation of virus-infected cells. We initially performed a three-dimensional (3D) analysis of SARS-CoV-2-infected cells using scanning electron microscopy (SEM) and TEM. By employing immunostaining and gene knockdown techniques, we provide compelling evidence that coatomer complex I (COPI) plays a crucial role in the transport of SARS-CoV-2 virions from the ERGIC. Our findings provide new insights into a previously unrecognized function of COPI in SARS-CoV-2 replication.

## Results

### Three-dimensional ultrastructural analysis of SARS-CoV-2 infected cells

To investigate the growth kinetics of SARS-CoV-2, we infected VeroE6/TMPRSS2 cells with the virus at a multiplicity of infection (MOI) of 1 and monitored viral titers in the supernatants up to 24 h post-infection (hpi) using a 50% tissue culture infectious dose (TCID_50_) assay. Concurrently, we assessed the expression of the SARS-CoV-2 N protein and double-stranded RNA (dsRNA) as a marker for viral genome replication over time using immunofluorescence assays (IFA). Notably, viral replication became detectable at 6 hpi, with titers steadily increasing up to 24 hpi (Figure 1a). In parallel, the expression of the N protein and dsRNA was observed in several cells at 6 hpi, with a subsequent increase in the number of virus-infected cells (Figure 1b). Our examination using ultrathin-section TEM revealed the presence of single virions within vesicular and tubular structures, which morphologically corresponded to the endoplasmic reticulum-Golgi intermediate compartment (ERGIC) at 6 and 12 hpi, while multiple virion-containing vacuoles were conspicuous at 24 hpi (Figure 1c). Therefore, for subsequent ultrastructural analysis, we employed samples collected at 24 hpi. We then embarked on a study of intracellular virion transport for release. Using SEM array tomography, we conducted a three-dimensional assessment of virion distribution within virus-infected cells (Supplemental Movie 1). This analysis unveiled various structures, including single virion-containing small vesicles, multiple virion-containing large vacuoles, and multiple virion-containing lysosomes dispersed within the cell, alongside numerous DMVs (Figure 1d). Virions were also observed within the membranes of the ERGIC (Figure 1e and Supplemental Movie 2) and large vacuoles during the budding process, which is consistent with prior reports (Figure 1f and Supplemental Movie 3) (10). To investigate how the progeny virions are released into the extracellular space, we focused on the membranous structures underneath the plasma membrane. Ultrathin-section TEM revealed small pits in the plasma membrane, on which the virions were present (Figure 2a). SEM array tomography affirmed that single virion-containing small vesicles were located beneath the plasma membrane (Figure 2b) and connected to the plasma membrane (Figure 2c) (Supplemental Movie 4). Notably, detailed ultrastructural analysis through TEM electron tomography revealed that the small vesicles/pits beneath the plasma membrane were coated with proteins (Figure 2d, e, and Supplementary Movie 5), suggesting virion release via exocytosis through these coated vesicles. Furthermore, ultrathin-section TEM exposed multiple virion-containing pleomorphic vacuoles near the plasma membrane, exhibiting several membrane protrusions forming a crown-like structure (Figure 2f). SEM array tomography corroborated these findings, suggesting that pleomorphic vacuoles with membrane protrusions are connected to the plasma membrane (Figure 2g and Supplemental Movie 6). TEM electron tomography revealed that the membrane protrusions on the pleomorphic vacuoles were also coated with proteins (Figure 2h, i, and Supplementary Movie 7), which is consistent with the observations in single virion-containing vesicles (Figure 2d, e). These findings suggest that virions are released via exocytosis through large pleomorphic vacuoles with coated membrane protrusions. Coated membrane protrusions were also observed in virion-containing lysosomes fused to the plasma membrane (Figure 2j and Supplemental Movie 8). In summary, our results indicate that while SARS-CoV-2 employs various vesicle types for virion transport, membrane-bound coat proteins likely play a crucial role in the vesicular transport of SARS-CoV-2 virions for their release.

**Figure 1.**
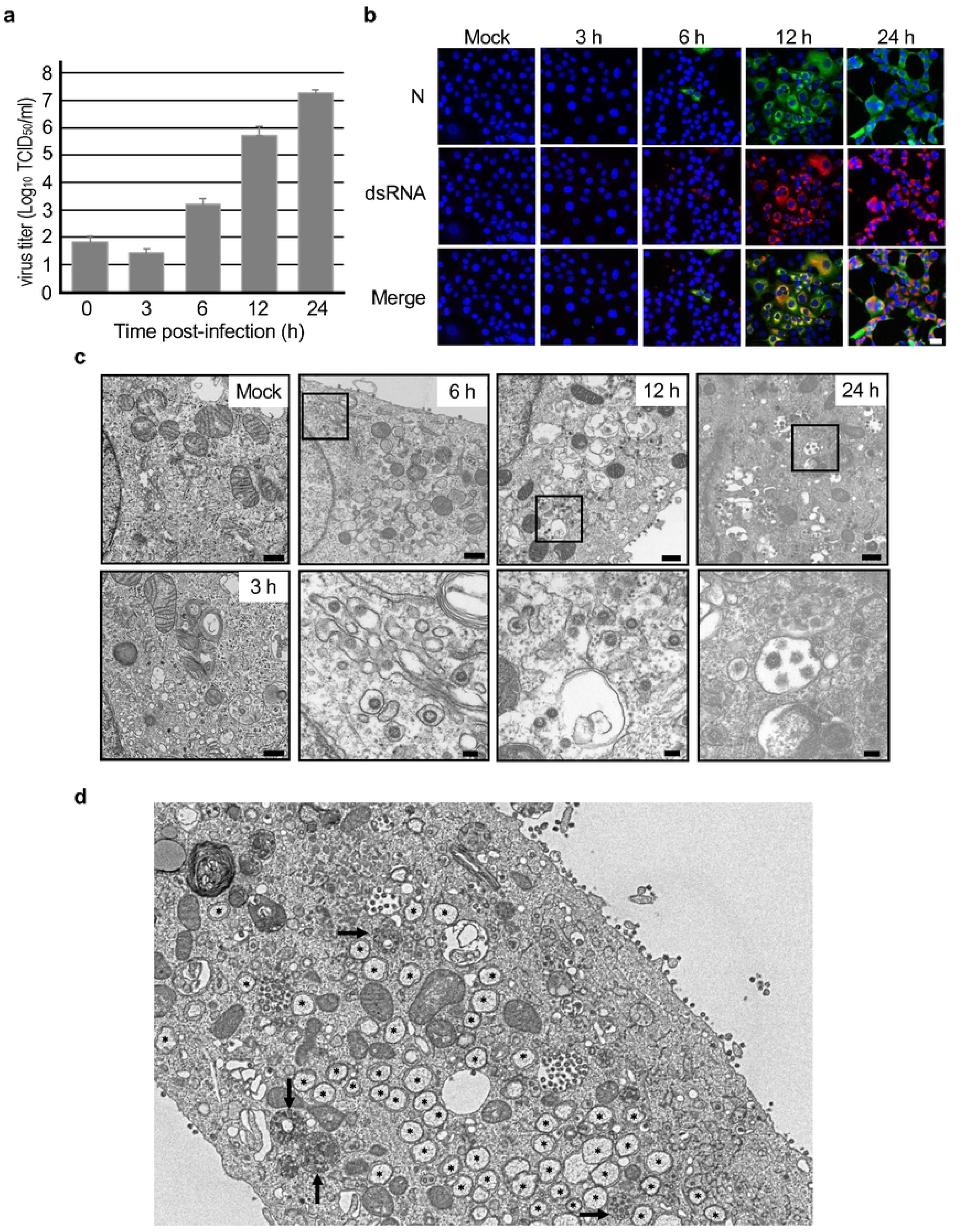
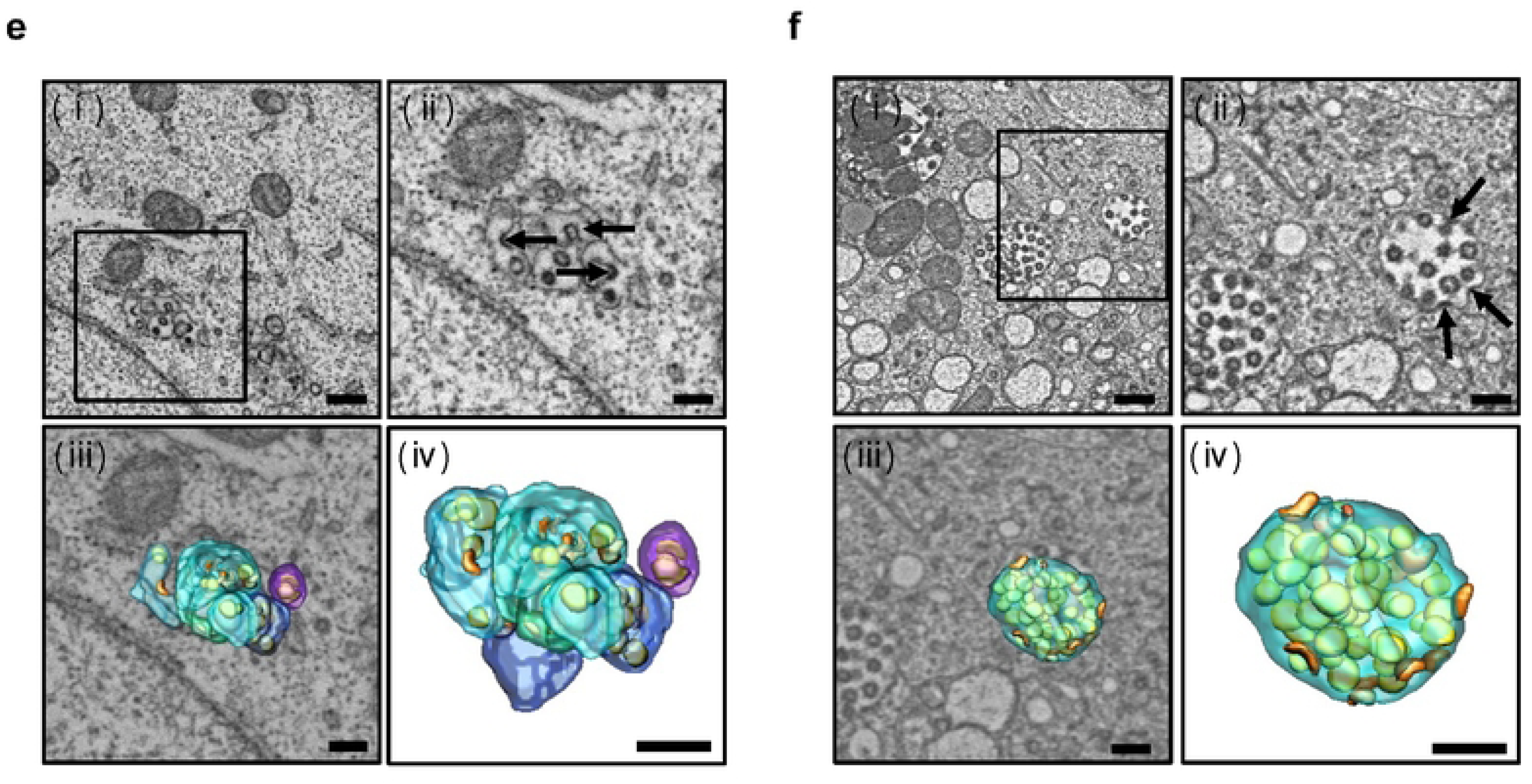
Replication kinetics of SARS-CoV-2 in VeroE6/TMPRSS2 cells. (a) Replication of SARS-CoV-2 in VeroE6/TMPRSS2 cells as determined by the 50% tissue culture infectious dose (TCID_50_) assay. The experiment was performed in biological triplicate. The error bars indicate the standard deviation (SD) of the three biological replicates. (b) Immunostaining images of mock- and SARS-CoV-2-infected VeroE6/TMPRSS2 cells using antibodies against the SARS-CoV-2 N protein (green) and double-stranded RNA (dsRNA) (red). Nuclei are stained with Hoechst 33342 (blue). The cells were infected with the virus at a multiplicity of infection (MOI) of 2 to collect the samples at 3 and 6 h post-infection (hpi) and at an MOI of 1 to collect samples at 12 and 24 hpi. Scale bar: 30 nm. (c) Ultrathin section images of SARS-CoV-2-infected VeroE6/TMPRSS2 cells at the indicated time points after infection. Box areas in images at 6, 12, and 24 hpi were magnified, respectively. Scale bars: 500 nm for low magnification images, and 100 nm for high magnification images. (d, e and f) Ultrastructure of endoplasmic reticulum-Golgi intermediate compartment (ERGIC) (e) and virion-containing vacuoles (f) in a virus-infected cell (d) imaged by scanning electron microscopy (SEM) array tomography. In (d), ∗ indicates double-membrane vesicles, and arrows indicate lysosomes. In (e) and (f), (i) Low magnification SEM image and (ii) High magnification image of the box area in (i) are shown. Arrows in (e) and (f) indicate virus budding. (iii) 3D rendering of ERGIC vesicles/tubules (light blue, blue, and purple), virion-containing vacuoles (light green), budding virions (orange), and budded virions (yellow) was superimposed on the image shown in (ii). (iv) 3D rendering of ERGIC vesicles/tubules (light blue, blue, and purple), virion-containing vacuoles (light green), budding virions (orange), and budded virions (yellow) shown in (iii). Nuc: nucleus. Scale bar: 400 nm (i), 200 nm (ii∼iv).

**Figure 2.**
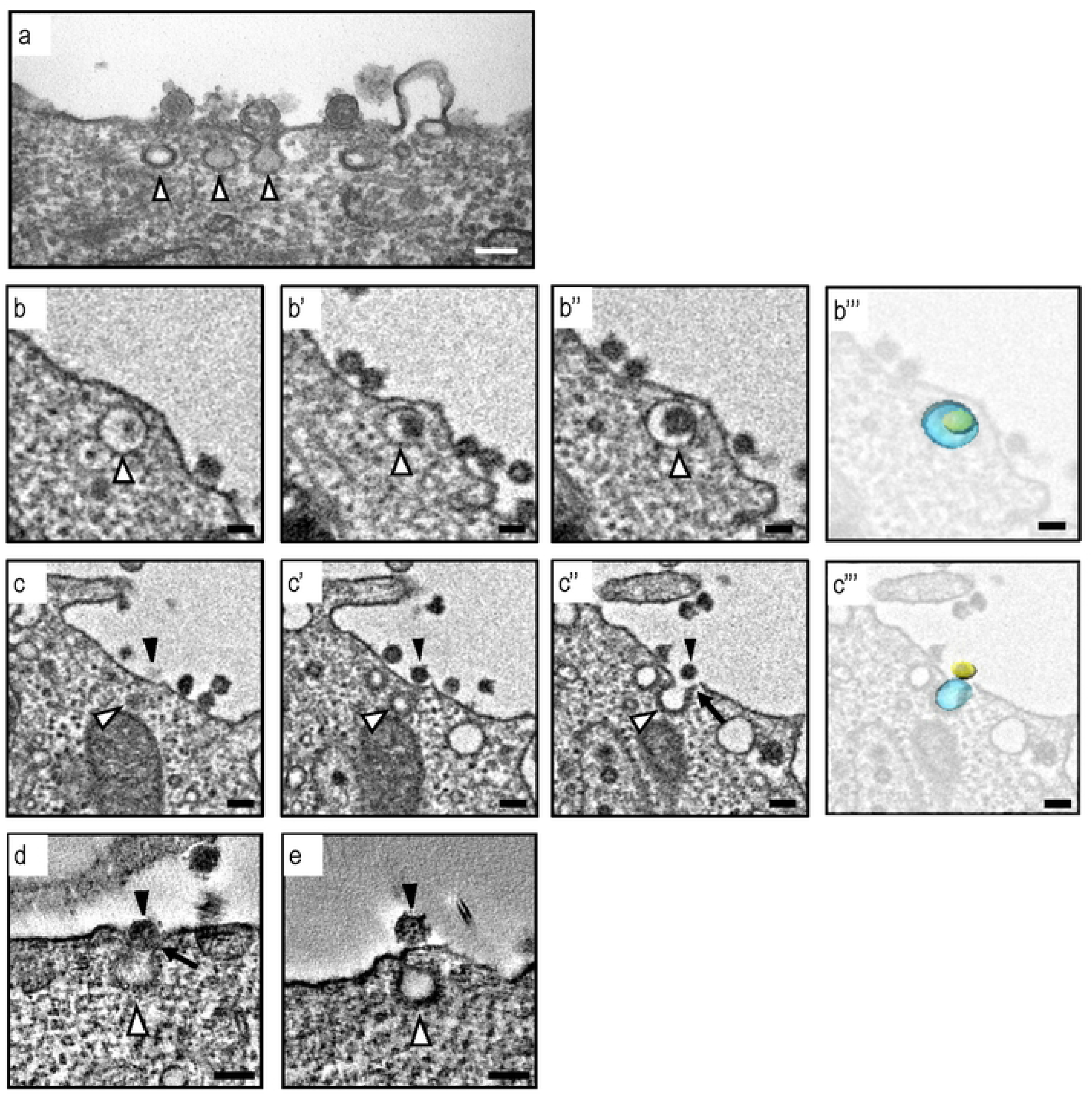
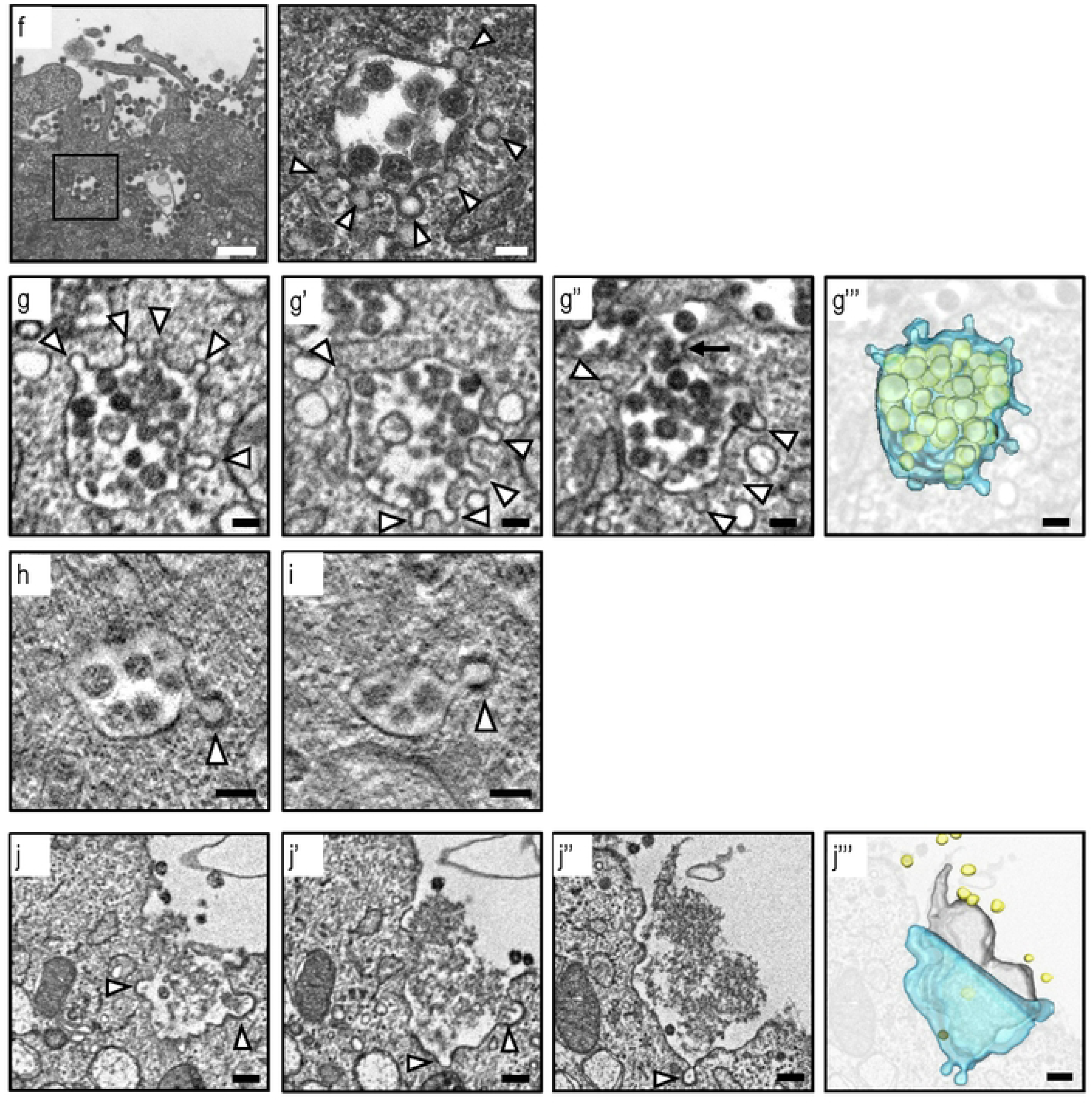
Ultrastructural analysis of SARS-CoV-2 virion transport using ultrathin-section TEM, SEM array tomography, and electron tomography. (a) An ultrathin-section image of the pits observed on the plasma membrane of a virus-infected cell. White arrowheads indicate pits close to SARS-CoV-2 virions. Scale bar: 100 nm. (b and c) Serial section images of the small vesicles containing single virions obtained using SEM array tomography. The b,b’, and b” and c, c’, and c” were taken from different sections of the same field, respectively. The b’” and c’” are superimposed images of b” and c” with 3D rendering of small vesicles (light blue) and virions (yellow), respectively. Black arrowheads indicate virions, and white arrowheads indicate small vesicle. Arrow indicates the membrane fusion between a vesicle and the plasma membrane. Scale bars: 100 nm. (d and e) Electron tomographic images of small vesicles transporting single virions. Black arrowheads indicate virions, and white arrowheads indicate coated proteins on the small vesicles. Arrow indicates the membrane fusion between a vesicle and the plasma membrane. Scale bars: 100 nm. (f) An ultrathin-section image of the multiple virions-containing large vacuoles near the plasma membrane. White arrowheads indicate membrane protrusion from the large vacuoles. Scale bars: 500 nm (left), 100 nm (right). (g) Serial section images of the large vacuole containing multiple virions obtained by SEM array tomography. The g, g’, and g” were taken from different sections of the same field. The g’” is a superimposed image of g” with 3D rendering of a large vacuole (light blue) and virions (yellow). White arrowheads indicate membrane protrusions. Scale bars: 100 nm. (h and i) Electron tomographic images of large vacuoles transporting multiple virions. White arrowheads indicate coated proteins on the membrane protrusions. Scale bars: 100 nm. (j) Serial section images of the virion-containing lysosome obtained by SEM array tomography. The j, j’, and j” were taken from different sections of the same field. The j’” is a superimposed image of j” with 3D rendering of the lysosome (light blue), virions (yellow), and degraded cellular components (gray). White arrowheads indicate membrane protrusions. Scale bars: 200 nm.

### Involvement of COPI in the vesicular transport of SARS-CoV-2 virions

It is well known that clathrin, COPI, and COPII are major coatomers that associate with membranous vesicles, regulating vesicular transport. Among these, COPI plays a critical role in protein-transporting vesicles responsible for retrograde transport from the ERGIC (14), the primary sites for SARS-CoV-2 assembly and budding (15). Hence, we postulated that the membrane-bound coat proteins found on vesicles and vacuoles containing virions are COPI, and COPI is involved in the transport of progeny virions from the ERGIC. To test this hypothesis, we examined the localization of β-COP or COPB2, subunits of the COPI complex, and ERGIC53, a specific ERGIC marker, in SARS-CoV-2 infected cells using IFA and immunoelectron microscopy (IEM). In mock-infected cells, COPB2 was primarily localized in the perinuclear region. However, during the later stages of SARS-CoV-2 infection, its distribution changed (Figure 3a). At 8 hpi, COPB2 colocalized with the SARS-CoV-2 S protein in the perinuclear region. By 16 hpi, both COPB2 and the SARS-CoV-2 S protein exhibited a punctate pattern throughout the cytoplasm (Figure 3a). In contrast to mock-infected cells, ERGIC53 exhibited a punctate distribution in the cytoplasm at 16 hpi and colocalized with the SARS-CoV-2 S protein (Figure 3b). Western blot analysis indicated that ERGIC53 expression in virus-infected cells was similar to that in mock-infected cells (Figure 3d), suggesting that SARS-CoV-2 replication may cause membrane alteration or fragmentation of the ERGIC. As reported previously, LAMP1, a lysosome marker, exhibited partial colocalization with the S protein (Figure 3c) (11). IEM revealed that both β-COP and ERGIC53 were present in virion-containing vesicles, small coated pits in the cytoplasm, and beneath the plasma membrane (Figure 3e, f). These findings collectively support the notion that COPI plays a role in intracellular virion transport through ERGIC-derived vesicles.

**Figure 3.**
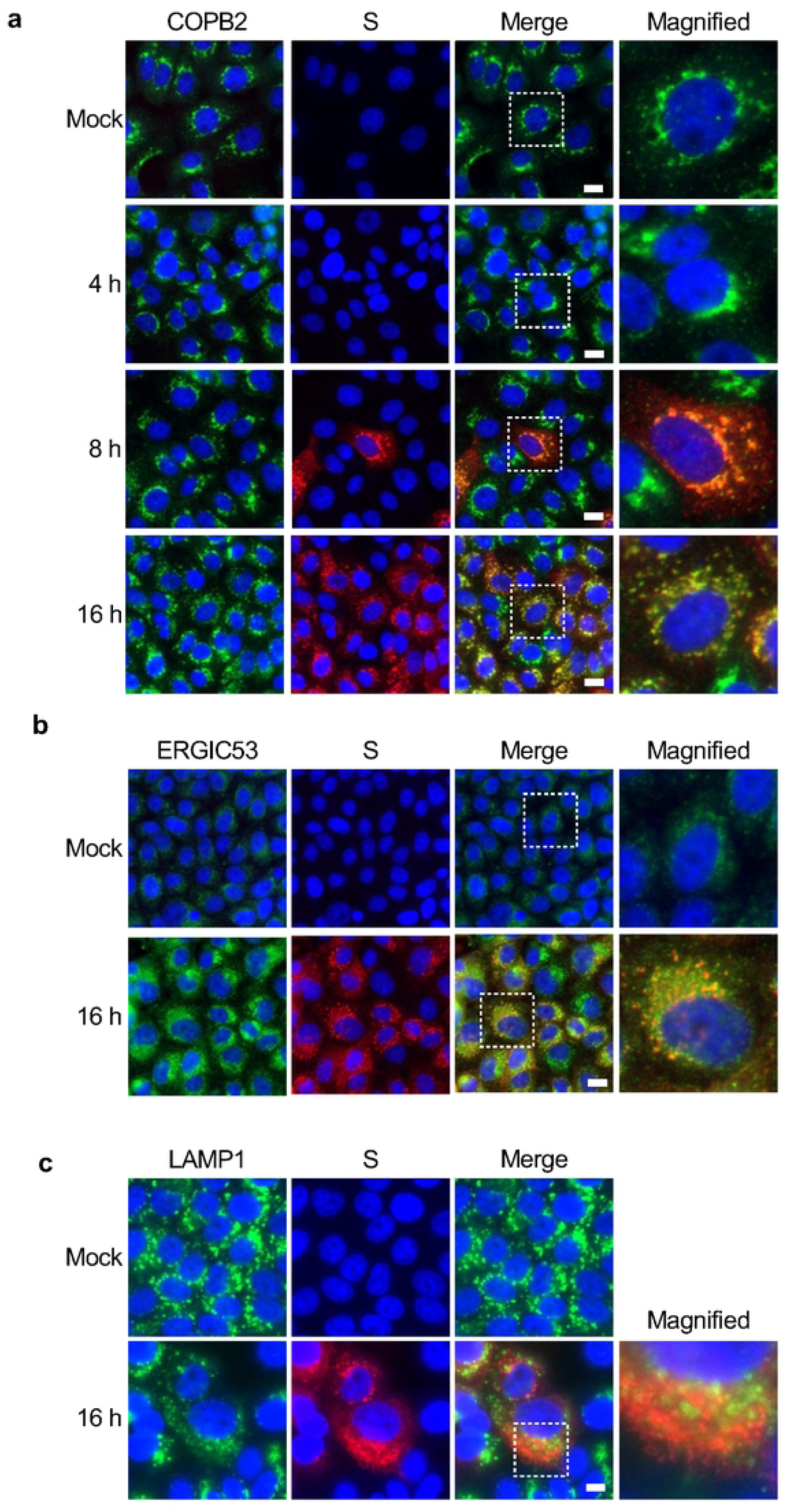
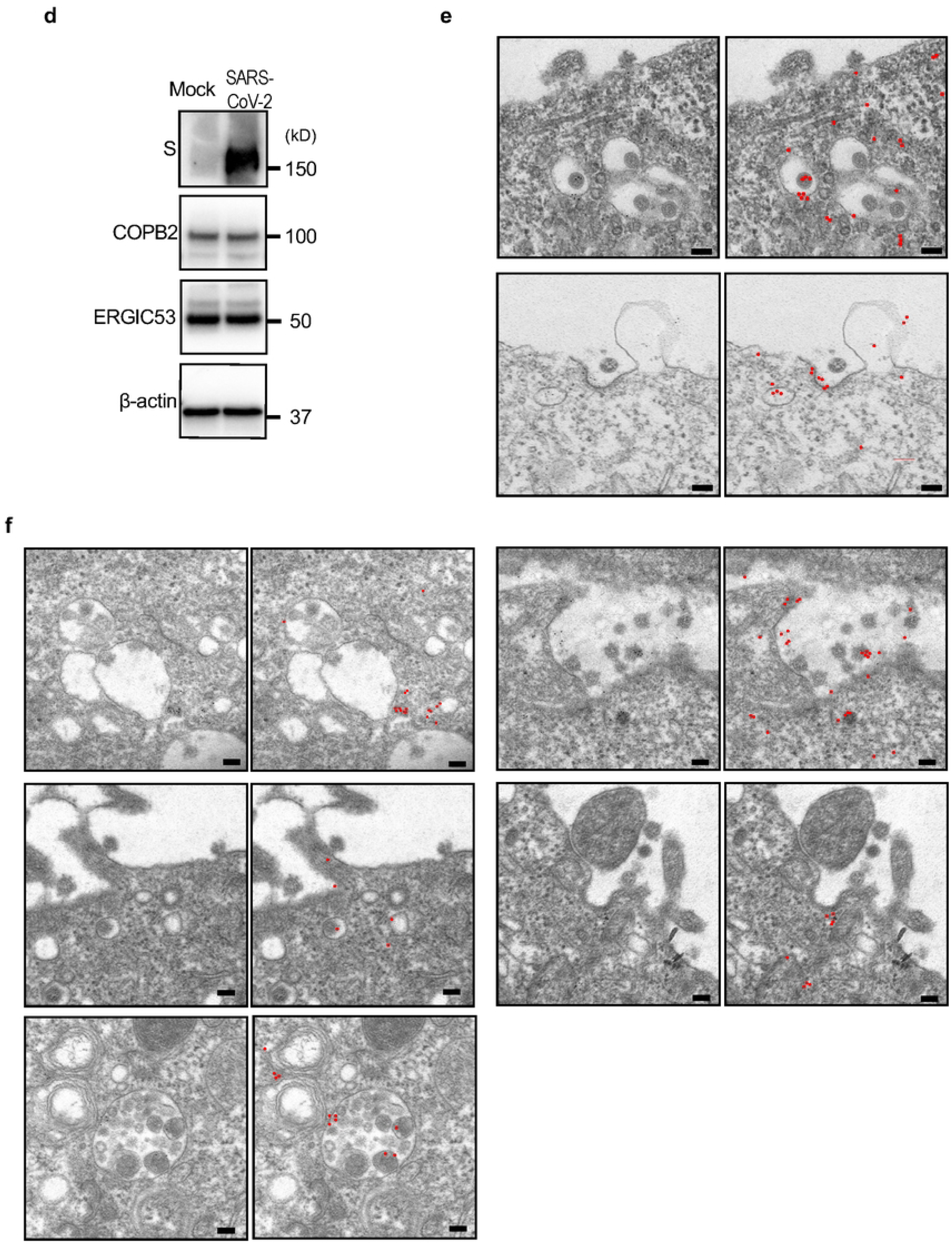
Intracellular localization of coatomer complex B2 (COPB2) and ERGIC53 in SARS-CoV-2-infected VeroE6/TMPRSS2 cells. (a, b, and c) VeroE6/TMPRSS2 cells were infected with SARS-CoV-2 at an MOI of 1, fixed at the indicated time points, and subjected to an immunofluorescence assay using antibodies against COPB2 (green in a), ERGIC53 (green in b), LAMP1 (green in c), and SARS-CoV-2 S protein (red). Nuclei were stained with Hoechst 33342 (blue). Box areas shown in merged images are magnified and displayed in the right-hand column. Scale bars: 10 µm. (d) Western blot analysis showing the expression levels of SARS-CoV-2 S protein, COPB2, ERGIC53, and β-actin in mock-infected and SARS-CoV-2-infected cells at 16 hpi. (e) Immunoelectron microscopy of SARS-CoV-2-infected cells using antibodies against β-COP, followed by secondary antibody conjugated with 6-nm gold beads. (right images) For clarity, gold beads were labeled with red and are shown in the right column. Scale bars: 100 nm. (f) Immunoelectron microscopy of SARS-CoV-2-infected cells using antibodies against ERGIC53, followed by secondary antibody conjugated with 6-nm gold beads. For clarity, gold beads were labeled with red and are shown on the right side. Scale bars: 100 nm.

### COPI facilitates the release of virion-containing vesicles from the ERGIC

To evaluate the significance of COPI in SARS-CoV-2 virion transport, we downregulated COPB2 expression using COPB2-targeting siRNAs. We confirmed a 50% reduction in COPB2 expression in COPB2-knockdown (COPB2-KD) cells compared to control cells without affecting cell viability (Figure 4a, b). COPB2 depletion significantly reduced SARS-CoV-2 replication to 2.5% at 24 hpi (Figure 4c), underscoring the essential role of COPI in efficient SARS-CoV-2 replication. To assess the impact of COPI on the vesicular transport of SARS-CoV-2 virions, we examined the intracellular localization of viral proteins in COPB2-KD cells using IFA. In control cells at 16 hpi, the SARS-CoV-2 S and M proteins, both of which are structural proteins of the virion, exhibited colocalization (Figure 4d) and were broadly distributed throughout the cytoplasm. In contrast, COPB2-KD cells exhibited altered distribution patterns for these viral proteins, with significantly increased signals in the perinuclear region (Figure 4e, e’). Additionally, the localization of ERGIC53 in COPB2-KD cells was constrained to the perinuclear region, where the SARS-CoV-2 S and M proteins colocalized, indicating the accumulation of virions in this region (Figure 4f). Ultrathin-section TEM revealed that ERGIC-like vacuoles containing virions were mainly limited to the perinuclear region in COPB2-KD cells, whereas virion-containing vesicles were dispersed throughout the cytoplasm in control cells at the same time point (Figure 4g). This suggests that the depletion of COPB2 impairs the transport of virion-containing vesicles from the ERGIC. Lastly, to gauge the impact of suppressing COPI vesicular transport on extracellular virion release, we examined the amounts of viral S, N, and M proteins in the cells and released into the supernatant at 8 hpi, which corresponds to a single replication cycle of the virus (Figure 4h). Additionally, we examined virus titers both within cells and in the supernatant at the same time point (Figure 4i). Although COPB2-KD cells exhibited comparable viral protein levels to control cells, the relative amounts of viral protein in the supernatants of COPB2-KD cells were significantly reduced to 15–25% compared to control cells (Figure 4h). Consistently, the virus titer detected in the supernatant of COPB2-KD cells was reduced to approximately 11% compared to that in the supernatant of control cells (Figure 4i). These results confirm the necessity of COPI vesicular transport for virion release from infected cells. Notably, the virus titer within COPB2-KD cells was significantly lower than that within control cells (Figure 4i), suggesting that COPB2 depletion hampers the proper assembly and formation of infectious virions. In summary, these findings underscore the pivotal role of COPI in facilitating the transport of virions from the ERGIC, thereby enabling their release from infected cells.

**Figure 4.**
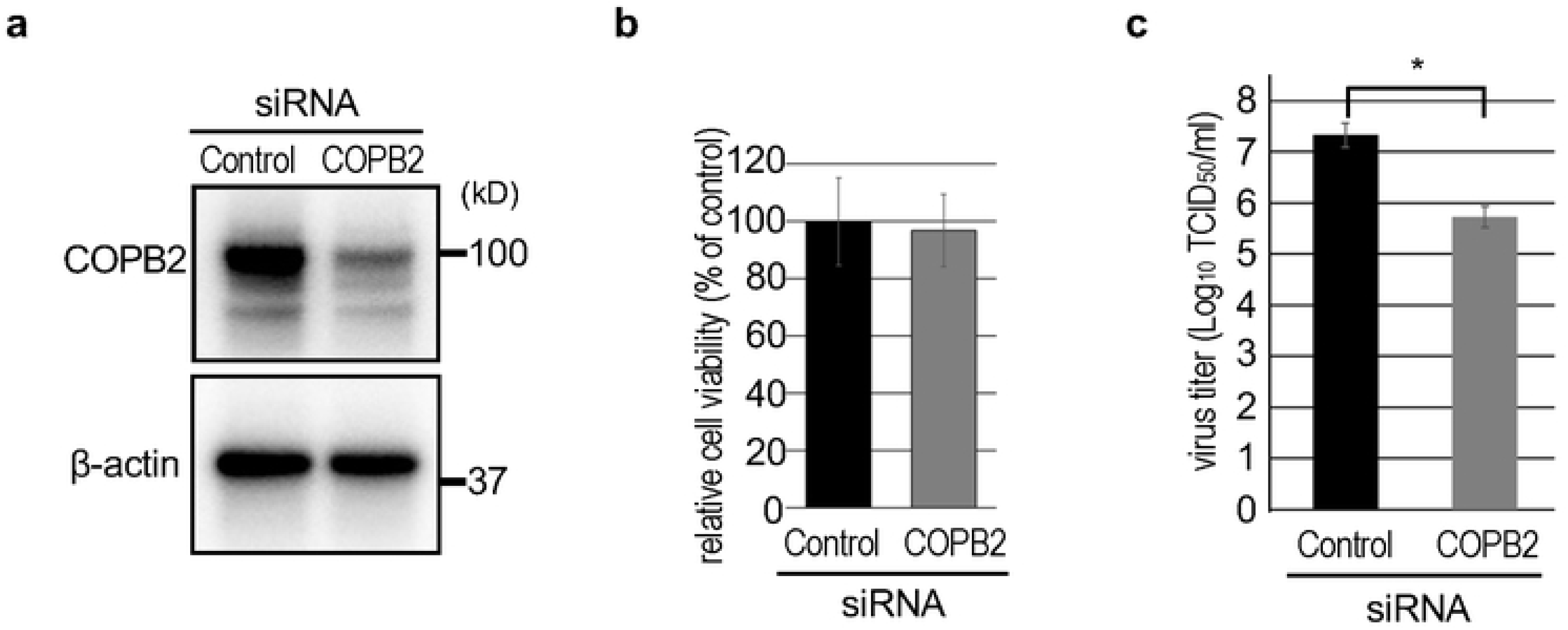
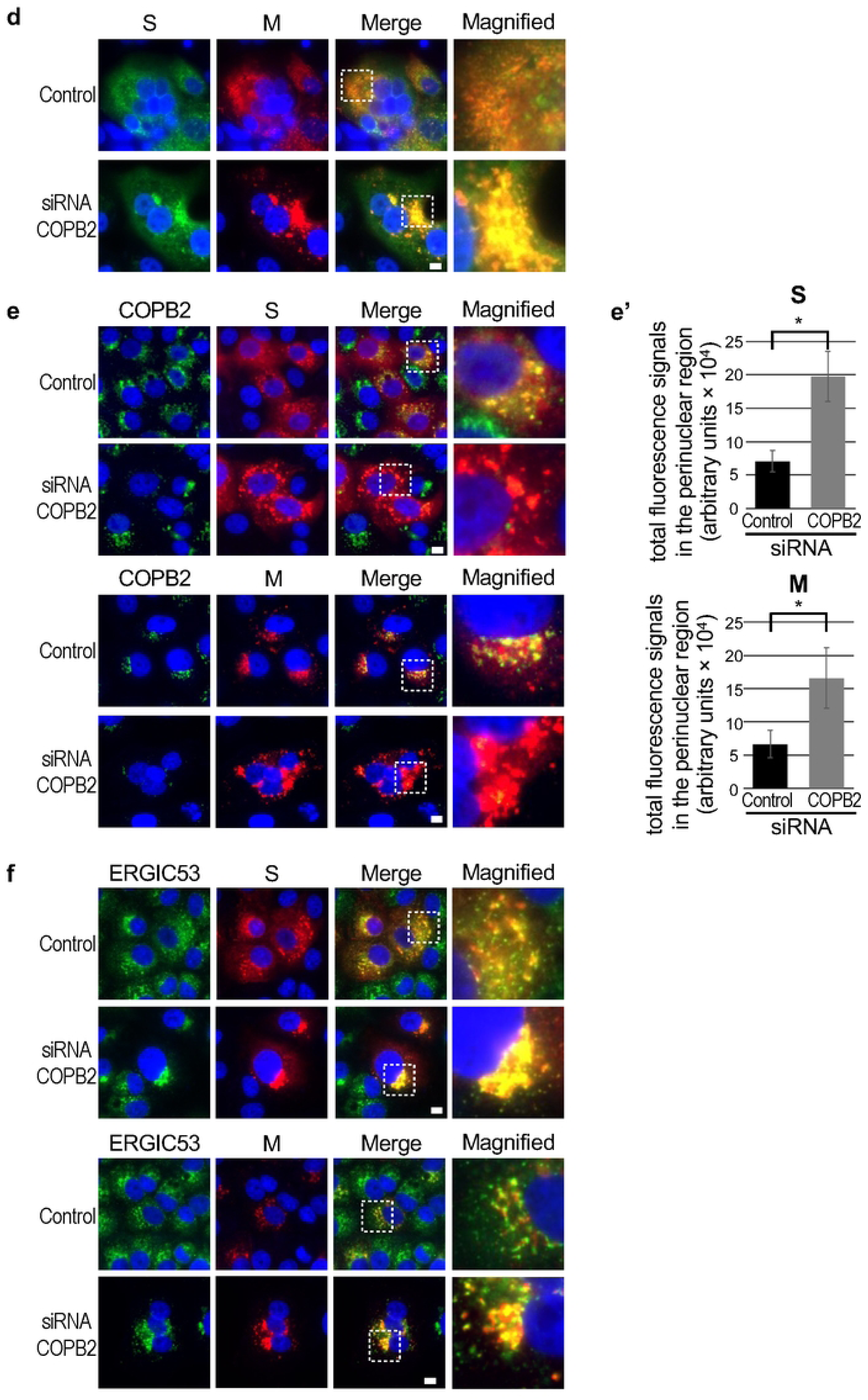
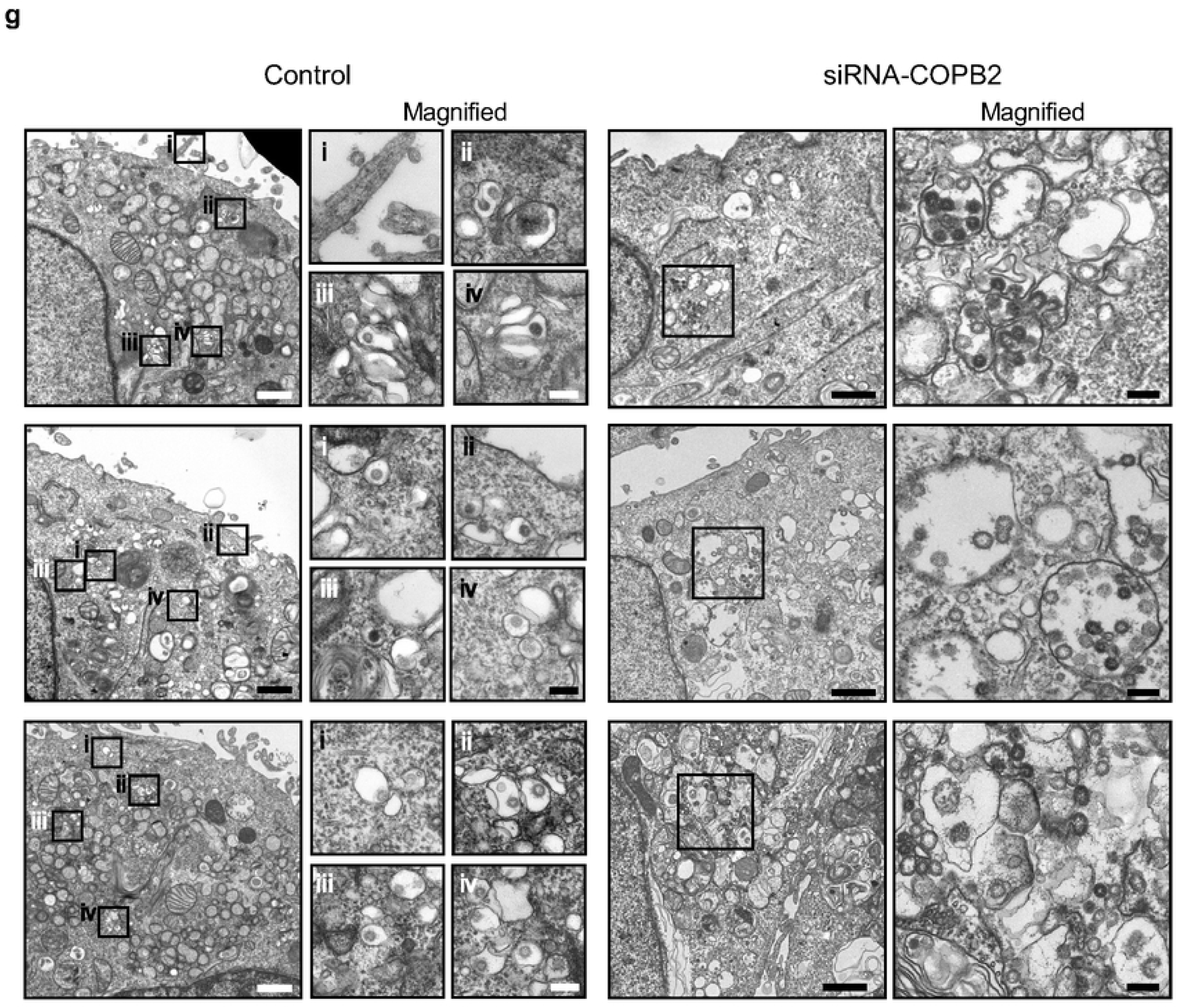
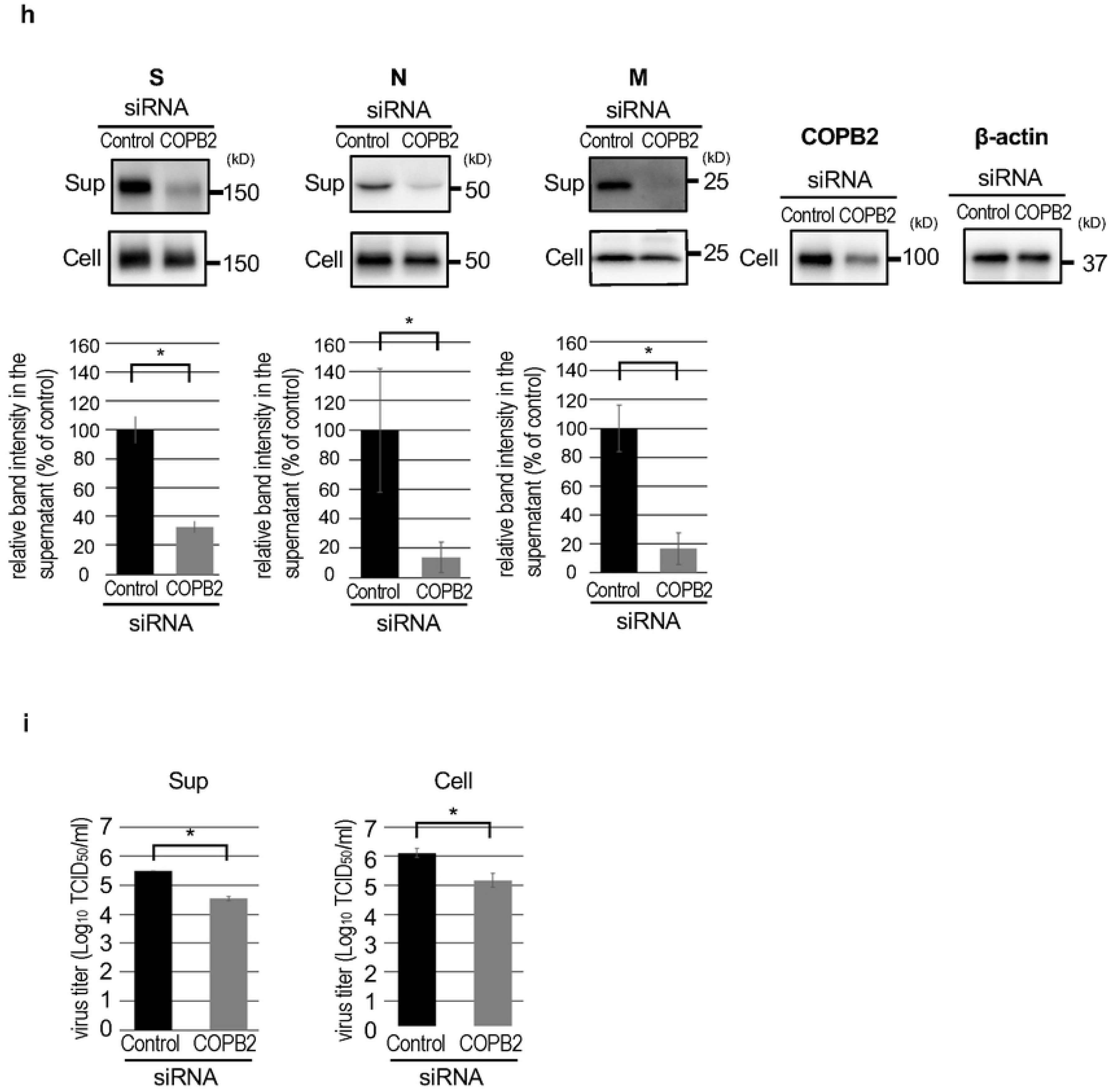
Impact of COPB2 depletion on SARS-CoV-2 replication. (a) The expression level of COPB2 in siRNA-treated Vero E6 cells was examined at 24 hours post-transfection by western blotting. (b) Cell viability was evaluated using a CellTiter-Glo assay at 12 hours post-transfection. Error bars indicate the standard deviation (SD) of three biological replicates. (c) Virus titers in the supernatants of COPB2-KD and control cells at 24 hpi. Error bars indicate the SD of three biological replicates. A two-tailed Student’s *t*-test was used to compare the two groups. \**p* < 0.05. (d,e,and f) Vero E6 cells were transfected with a control siRNA or siRNA against COPB2, infected by SARS-CoV-2, and fixed at 16 hpi. COPB2 (green in e), ERGIC53 (green in f), SARS-COV-2 S protein (green in d, red in e and f), and SARS-CoV-2 M protein (red in d and f) were immunolabeled. Nuclei were stained with Hoechst 33342 (blue). Bars:10 µm. (e’) The graphs display total intensities of S and M proteins around the nuclei of virus-infected control and COPB2-KD cells. Twenty virus-infected cells were randomly chosen, and the arbitrary units were calculated by the sum of the multiplication of each intensity by the areas. Error bars indicate the SD of three biological replicates. A two-tailed Student’s *t*-test was used to compare the two groups. \**p* < 0.05. (g) Ultrathin-section TEM images of control and COPB2-KD cells infected with SARS-CoV-2 at 16 hpi. Control: Respective box regions (i-iv) are magnified and shown on the right. Scale bars: 1 µm (low magnification images) and 100 nm (high magnification images). siRNA-COPB2: Box regions are magnified and shown on the right. Scale bars: 1 µm (low magnification images) and 100 nm (high magnification images). (h) Detection of viral structural proteins in virus-infected control and COPB2-KD cells by western blot analysis. The experiment was performed in triplicate, and the band intensities of respective viral proteins were quantified and are shown as respective graphs. Error bars indicate the SD of three biological replicates. A two-tailed Student’s *t*-test was used to compare the two groups. \**p* < 0.05. (i) Virus titers in the supernatants and cells of COPB2-KD and control cells at 8 hpi. Error bars indicate the SD of three biological replicates. A two-tailed Student’s *t*-test was used to compare the two groups. \**p* < 0.05.

## Discussion

Progeny virions of SARS-CoV-2, which bud into the lumen of the ERGIC, necessitate transport through membranous structures to exit the cell. Yet, the molecular mechanisms governing this process have remained elusive. In this study, we conducted a three-dimensional ultrastructural analysis of virus-infected cells using SEM array tomography and electron tomography and discovered that the vesicles containing virions were outfitted with membrane coat proteins. Immunostaining revealed that the coat protein in question was COPI, and these virion-containing vesicles originated from the ERGIC. Crucially, COPI depletion hindered the transport of virions from the ERGIC, resulting in an accumulation of the progeny virions within the ERGIC, a notable reduction in virion release, and a decrease in viral replication. Collectively, our findings revealed the pivotal role of COPI during the SARS-CoV-2 life cycle.

During retrograde transport from the Golgi apparatus and ERGIC to the ER, the COPI complex plays a central role in cargo sorting, cargo uptake, vesicle formation, and vesicular release from membranes, giving rise to COPI-coated cargo-transporting vesicles (16). In a previous kinome-wide siRNA screening for SARS-CoV, COPI was identified as a proviral host factor, although its precise role remains unknown (17). Recent studies have also reported the utilization of the COPI complex for transporting post-translationally modified S proteins from the Golgi apparatus back to the ERGIC for virion assembly during SARS-CoV-2 replication (18, 19). In our investigation, we unearthed an additional role of COPI, one that involves the intracellular transport of progeny virions. It appears that COPI is indispensable for the formation of vesicles that contain virions on the ERGIC membrane, thereby facilitating the transport of progeny virions from the ERGIC. Intriguingly, although COPI coats typically dissociate from vesicles shortly after their release during retrograde transport, they continued to be observed on many vesicles containing virions, even after transport to the cell surface. Future studies should explore whether COPI associated with virion-containing vesicles plays additional roles in intracellular transport and/or the fusion process at the plasma membrane for virion release.

Because spherical virions were observed within the ERGIC in COPI-KD cells (Figure 4g), we expected that COPI depletion would not critically affect progeny virion formation. However, the reduced production of infectious virions within COPB2-KD cells compared to control cells (Figure 4i) suggests that appropriate assembly of viral structural proteins would be hindered due to the depletion of COPB2. This is likely caused by the inhibition of retrograde transport of the S protein from the Golgi apparatus to the ERGIC (18, 19), resulting in an insufficient amount of S protein being provided for virion assembly.

A recent study identified lysosomes as vesicular transporters of progeny virions (11). Additionally, it has been suggested that progeny virions are transported through single- (13) and multiple-virion-containing vesicles (12), although the origins of these vesicles remain unidentified. In our ultrastructural analysis, single-virion-containing vesicles were predominantly observed at 6 and 12 hpi, whereas multiple-virion-containing vesicles and virion-containing lysosomes were more frequent at 24 hpi. This suggests that these vesicular transport systems are not mutually exclusive and that the vesicles observed in infected cells depend on the timing of viral infection and replication. Thus, based on our findings and in conjunction with previous reports, we propose a hypothetical model where the vesicular transport of virions from the ERGIC is mediated by COPI (Figure 5). During the early stages of viral infection, the efficiency of progeny virion formation is relatively low due to the reduced expression of viral proteins and the viral genome. Consequently, a single virion buds into the lumen of the ERGIC, and subsequently, single virion-containing vesicles are released from the ERGIC, which is mediated by the COPI complex (Figure 5a). These single virion-containing vesicles are then individually transported to the cell surface. As viral expression levels increase during the later stages of infection, the efficiency of progeny virion formation also increases, resulting in the budding of multiple virions into the ERGIC lumen. Consequently, many single-virion-containing vesicles are efficiently and simultaneously released from the ERGIC by the COPI complex, which may lead to the fusion of these vesicles to form larger vesicles during the transport of virions to the cell surface (Figure 5b). Alternatively, multiple virion-containing ERGIC vesicles may be released from the ERGIC, although it is unclear whether the COPI complex has the ability to form large vesicles containing multiple virions. Our analysis did not reveal budding virions at the lysosomal membrane; thus, it is presumed that lysosomes may fuse with virion-containing ERGIC-derived vesicles during their transport to the cell surface (Figure 5c). Therefore, we propose that regardless of their size, virion-containing vesicles originate from the ERGIC, and their transport from the ERGIC is regulated by the COPI complex.

**Figure 5.**
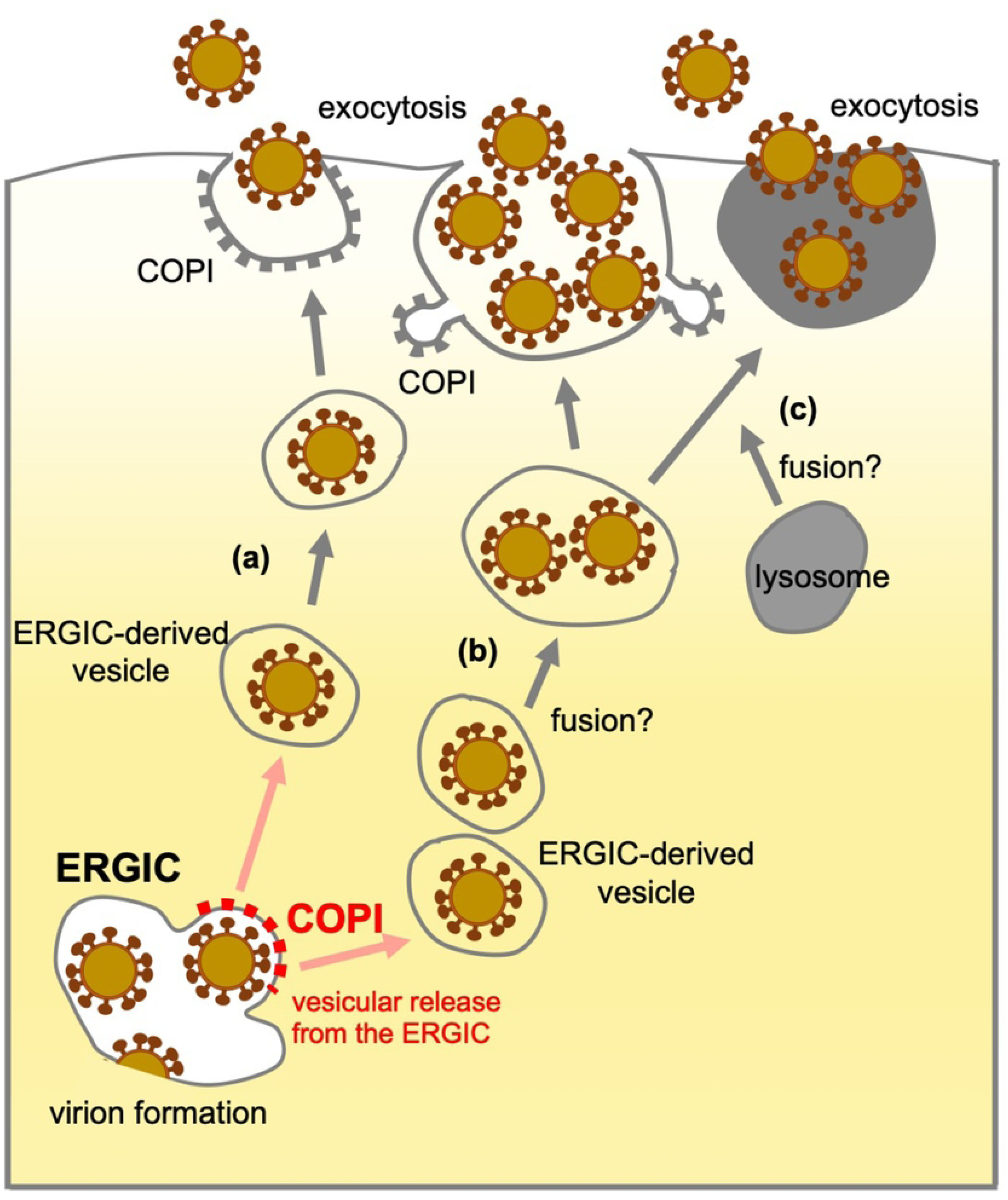
A proposed model of the vesicular transport of SARS-CoV-2. Virions bud into the lumen of the ERGIC. (a) Thereafter, single virion-containing vesicles are released from the ERGIC, which is mediated by the COPI complex. (b) Many single virion-containing vesicles are released from the ERGIC as the budding efficiency of virions increases during the later stages of infection. During their transport to the cell surface, multiple virion-containing vesicles may be formed by the repetitive fusion of virion-containing vesicles. (c) Virion-containing vesicles may be fused with lysosomes during their transport to the cell surface. Red dots on the ERGIC represent COPI.

In summary, our study has demonstrated that COPI plays a critical role in the vesicular transport of SARS-CoV-2 virions from the ERGIC. This work unveils a novel role of COPI in the SARS-CoV-2 life cycle and underscores its significance in replication. Given its involvement in multiple steps, including virion transport from the ERGIC, retrograde transport of the S protein during viral replication, and proper assembly of infectious virions, COPI represents a potential target for novel antiviral development.

## Materials and Methods

### Cells and virus

Vero E6 and VeroE6/TMPRSS2 cells (20) were maintained in Dulbecco’s modified Eagle’s medium (DMEM) containing 10% fetal calf serum at 37 °C under 5% CO_2_. The SARS-CoV-2 isolate (SARS-CoV-2/Hu/DP/Kng/19-027), provided by the Kanagawa Prefectural Institute of Public Health, was propagated in VeroE6/TMPRSS2 cells, and titers were determined by TCID_50_. All SARS-CoV-2 experiments were performed in a biosafety Level 3 containment laboratory at the Institute for Life and Medical Sciences, Kyoto University.

### Antibodies

The commercially available primary antibodies used for immunofluorescence, western blotting, and immuno-electron microscopy were as follows: anti-dsRNA mouse monoclonal antibody J2 (10010200; Scicons; Nordic MUbio, Susteren, Netherlands), anti-β-COP rabbit polyclonal antibody (ab2899; abcam, UK), anti-COPB2 rabbit polyclonal antibody (A304-523A; Bethyl, Montogomery, TX, USA), (ab192924; abcam, UK), anti-SARS-CoV-2-S rabbit polyclonal antibody (NB100-56578; Novus Bililigicals, Centennial, CO, USA), anti-SARS-CoV-2-N rabbit polyclonal antibody (GTX135357; GeneTex, Irvine, CA, USA), anti-SARS-CoV-2-N mouse monoclonal antibody (ZMS1075; Merck, Darmstadt, Germany), anti-LAMP1 rabbit polyclonal antibody (9091T; Cell Signaling Technology, Danvers, MA, USA), and anti-β-actin mouse monoclonal antibody (ab8226; abcam, UK). In-house anti-SARS-CoV-2-S mouse monoclonal antibody (#7), anti-SARS-CoV-2-M mouse monoclonal antibodies (YN7730-01), and mouse antiserum against SARS-CoV-2 M (#27-1-3) were generated by immunizing mice with the purified receptor-binding domains of the SARS-CoV-2 S and SARS-CoV-2-M proteins, respectively. The secondary antibodies used were Alexa Fluor 488-conjugated anti-rabbit (A11008; Thermo Fisher Scientific, Waltham, Ma, USA), Alexa Fluor 555-conjugated anti-mouse (A21422; Thermo Fisher Scientific), horseradish peroxidase-conjugated anti-mouse (NA931; GE Healthcare, Chicago, IL), anti-rabbit (NA934; GE Healthcare), and 6-nm gold-conjugated anti-rabbit (711-195-152; Jackson ImmunoResearch) antibody (711-195-152; Jackson ImmunoResearch, West Grove, PA, USA).

### Virus titration

The viruses were titrated using the TCID_50_ assay, as described previously (21). Briefly, VeroE6/TMPRSS2 cells were seeded in a 96-well plate one day prior to the assay. Serially diluted virus samples were inoculated into the cells and incubated at 37 °C for 4 days. Thereafter, the cells were observed under a microscope to assess cytopathic effects. The viral titer (TCID_50_/ml) was calculated using the Reed–Muench method (22). For the titration of virus titer within cells, VeroE6 or COPI-KD cells infected with SARS-CoV-2 at an MOI of 10 were washed with a medium at 8 hpi, fresh medium was added, and then cells were freeze-thawed three times to break the infected cells. After removing debris by centrifugation, the supernatant containing the virus was collected and subjected to titration using the TCID_50_ assay.

### Immunofluorescence assay

VeroE6 or VeroE6/TMPRSS2 cells infected with SARS-CoV-2 were fixed with 4% paraformaldehyde at 4 °C for 1 h and permeabilized with 0.1% Triton X-100 for 10 min. After blocking with Blocking One solution (Nacalai Tesque) for 30 min, the cells were treated with antibodies against the SARS-CoV-2 N protein (1:1000 dilution, GTX135357, ZMS1075), dsRNA (1:500 dilution, 10010200), COPB2 (1:1000 dilution, ab192924), SARS-CoV-2-S (1:200 dilution, #7), (1:100 dilution, NB100-56578), or SARS-CoV-2-M (1:100 dilution, YN7730-01), followed by secondary antibodies against rabbit and mouse IgG. Nuclei were stained with Hoechst 33342 (1:1000 dilution, H3570, Thermo Fisher Scientific). Fluorescent images were obtained using a BZ-X800 fluorescence microscope (Keyence, Japan). For quantification of the total fluorescence signals, the signals corresponding to S and M proteins around the periphery of the nuclei were automatically examined. Initially, a mask was created by extending about 4 µm from the nuclear boundary of each cell, followed by a smoothing process applied to the mask. Then, the nuclear area was subtracted from the masked area to quantify the signals only around the nuclear periphery. The total signal within the mask was calculated by multiplying the signal area of viral proteins by the intensities within the masked area. All data was analyzed using BZ-X800 Analyzer (Keyence, Japan).

### Sample preparation for electron microscopy

Vero E6 or VeroE6/TMPRSS2 cells infected with SARS-CoV-2 were fixed with 2.5% glutaraldehyde in 0.1M cacodylate buffer on ice for 1 h and postfixed with 1% osmium tetroxide on ice for 1 h. The fixed samples were dehydrated using a series of ethanol gradients, substituted with propylene oxide, and embedded in epoxy resin.

### Ultrathin-section TEM, electron tomography, and immunoelectron microscopy

The epoxy blocks were trimmed, and ultrathin sections (∼50 nm) were cut using an ultramicrotome (EM UC7; Leica, Wetzlar, Germany) equipped with an ultra-diamond knife (Diatome, Biel, Switzerland). For electron tomography, ultrathin sections (∼200 nm) were cut, stained with 2% uranyl acetate and Reynold’s lead, and coated with carbon on both sides. The tilt series were recorded with a 200-kV TEM (TalosF200C, FEI, Thermo Fisher Scientific), and the tomograms were reconstructed using the simultaneous iterative reconstruction technique using IMOD (23). For immunoelectron microscopy, ultrathin sections were prepared on nickel grids, as described above, and incubated with saturated sodium periodate solution, followed by 0.2 M glycine in phosphate buffered saline (PBS) buffer (Noda et al., Nature 2006). After washing with PBS, sections were incubated with blocking buffer and anti-β-COP antibodies (1:100 dilution, ab2899). Thereafter, sections were washed with PBS and incubated with anti-rabbit immunoglobulin conjugated to 6-nm gold particles (1:30 dilution, 711-195-152). Ultrathin sections were stained with uranyl acetate and lead citrate and observed in an HT-7700 (Hitachi High-Tech Corporation, Japan) at 80 kV.

### SEM array tomography

The epoxy blocks were trimmed, and serial sections (30-nm thick) were cut with an ultramicrotome (EM UC7) using an ultra-diamond knife (Diatome). Sections were mounted on silicon wafers. Serial sections were stained with uranyl acetate and lead citrate and coated with osmium coater (HPC-SW; Vacuum Device Co.). Images were obtained using field-emission SEM equipped with an auto capture for array tomography system (Regulus8220 ACAT; Hitachi High-Tech Corporation, Tokyo, Japan) with a backscattered electron detector at 2 kV. Three-dimensional image reconstruction was performed using Image-Pro software (Hakuto, Tokyo, Japan).

### siRNA treatment

Three siRNAs targeting COPB2 (# 9276-1, # 9276-2, # 9276-3) were purchased from Bioneer Corporation (USA). AllStars Negative Control siRNA (# 1027280; GIAGEN, Hilden, Germany) was used as a negative control. Vero E6 cells cultured in 24-well plates were transfected with a mixture of 25 nM siRNAs using the IT-X2 reagent (Mirus Bio LLC, USA). A second transfection was performed 24 h after the first transfection, which was subjected to viral infection 1 d after the second transfection. The downregulation of COPB2 was evaluated by western blotting.

### Cell viability assay

To evaluate cell viability, cells were collected 24 h after siRNA treatment, and the amount of intracellular ATP was measured using the CellTiter-Glo assay according to the manufacturer’s protocol (Promega, Madison, WI, USA).

### Western blotting

Virus-infected cells and the supernatants were dissolved with Tris-glycine SDS sample buffer (Thermo Fisher Scientific, Waltham, MA, USA), boiled for 5 min in the absence of reducing agent, and subjected to sodium dodecyl sulfate-polyacrylamide gel electrophoresis (SDS-PAGE). The proteins were electroblotted onto Immobilon-P transfer membranes (Merck). Thereafter, membranes were blocked with Blocking One for 30 min at room temperature and then incubated with primary antibodies, anti-COPB2 antibody (1:5000 dilution), anti-SARS-CoV-2 S protein (1:1000 dilution, NB100-56578), anti-SARS-CoV-2 N protein (1:10000, GTX135357), a mouse antiserum against SARS-CoV-2 M protein (1:1 dilution), or anti-β-actin antibody (1:10000 dilution) overnight at 4 °C. After incubation with horseradish peroxidase-conjugated secondary antibodies (1:10000 dilution) for 1 h at room temperature, the blots were developed using Chemi-Lumi One Super (Nacalai Tesque).

## Statistical analysis

Statistically significant differences in cell viability, virus titers, and the relative amounts of viral proteins were determined using a two-sided two-sample Student’s *t*-test. For viral titers, the statistical tests were performed on the log_10_ transformed data. In the figures, asterisks denote statistical significance as calculated by the Student’s *t*-test (\**p* < 0.05). The error bars indicate the standard deviation (SD).

## Acknowledgments

We would like to thank Rumi Sato and Takaharu Okada (RIKEN IMS) for offering the SEM array tomography system. We also thank Kehong Liu (Kyoto University) for her technical assistance in the production of the anti-M antibodies. This work was supported by the JSPS Core-to-Core Program A, Advanced Research Networks (JPJSCCA20190008 and JPJSCCA20240006), JSPS Grant-in-Aid for Challenging Exploratory Research (22K19431), JST Core Research for Evolutional Science and Technology [JPMJCR20HA], AMED (24fk0108694h0001, JP23fm0208101 and 22KK0115), Grant for Joint Research Project of the Institute of Medical Science at the University of Tokyo, Joint Usage/Research Center Program of the Institute for Life and Medical Sciences at Kyoto University, the Takeda Science Foundation (to TN), and AMED Basis for Supporting Innovative Drug Discovery and Life Science Research (BINDS, JP22ama121007) (to NN).

## Supplementary Figure legends

**Movie S1.** 3D image of a SARS-CoV-2-infected VeroE6/TMPRSS2 cell reconstructed using SEM array tomography.

**Movie S2.** 3D image of ERGICs containing progeny virions reconstructed using SEM array tomography. 3D rendering was performed for ERGIC vesicles/tubules (light blue, blue, and purple), budding virions (orange), and budded virions (yellow).

**Movie S3.** 3D image of a multiple virion-containing large vacuole reconstructed using SEM array tomography. 3D rendering was performed for virion-containing large vacuole (light green), budding virions (orange), and budded virions (yellow).

**Movie S4.** 3D image of single virion-containing small vesicles reconstructed using SEM array tomography. 3D rendering was performed for single virion-containing small vesicles (light blue), and budded virions (yellow).

**Movie S5.** Electron tomographic image of small vesicles/pits coated with proteins that are located beneath the plasma membrane of virus-infected cell.

**Movie S6.** 3D image of a multiple virions-containing large vacuole that is located beneath the plasma membrane of the virus-infected cell, reconstructed using SEM array tomography. 3D rendering was performed for virion-containing large vacuole (light blue), budding virions (orange), and budded virions (yellow).

**Movie S7.** Electron tomographic image of multiple virion-containing vesicles located beneath the plasma membrane of virus-infected cell. Some virion-containing vesicles have protrusion coated with proteins.

**Movie S8.** 3D image of a multiple virion-containing lysosome that is fused with the plasma membrane of virus-infected cell, reconstructed using SEM array tomography. 3D rendering was performed for virion-containing lysosome (light blue), degraded cellular components (gray), and budded virions (yellow).

